# Differential metabolism between biofilm and suspended *Pseudomonas aeruginosa* cultures in bovine synovial fluid by 2D NMR-based metabolomics

**DOI:** 10.1101/2022.06.16.496519

**Authors:** Abigail Leggett, Da-Wei Li, Lei Bruschweiler-Li, Anne Sullivan, Paul Stoodley, Rafael Brüschweiler

**Affiliations:** Ohio State Biochemistry Program, The Ohio State University, Columbus, OH, USA; Department of Chemistry and Biochemistry, The Ohio State University, Columbus, OH, USA; Department of Microbial Infection and Immunity, The Ohio State University, Columbus, OH, USA; Campus Chemical Instrument Center, The Ohio State University, Columbus, OH, USA; College of Medicine, Wexner Medical Center, Columbus, OH, USA; Department of Orthopaedics, The Ohio State University, Columbus, OH, USA; Department of Microbiology, The Ohio State University, Columbus, OH, USA; National Biofilm Innovation Centre (NBIC) and National Centre for Advanced Tribology at Southampton (nCATS), Mechanical Engineering, University of Southampton, Southampton, UK; Department of Biological Chemistry and Pharmacology, The Ohio State University, Columbus, OH, USA

**Author notes:** Corresponding author, (RB), (PS).

## Abstract

Total joint arthroplasty is a common surgical procedure resulting in improved quality of life; however, a leading cause of surgery failure is periprosthetic joint infection. Periprosthetic infection often involves biofilms, making treatment challenging. Periprosthetic joint infections are difficult to diagnose by traditional culturing methods and there are no clinical biomarkers for the presence of biofilms. Further, the metabolic state of pathogens in the joint space is difficult to diagnose, the mechanism of their tolerance to antibiotics and host defenses is not well understood, and their culturing in the laboratory is challenging. Thus, there is a critical need for improved pathogen- and phenotype-specific diagnosis as well as improved treatment strategies toward better patient outcomes. Here, we present a quantitative, untargeted NMR-based metabolomics strategy for *Pseudomonas aeruginosa* suspended culture and biofilm phenotypes grown in bovine synovial fluid. We identified 21 unique metabolites as potential markers of *P. aeruginosa* and one unique marker of the biofilm phenotype in synovial fluid. Significant differences in metabolic pathways were found between the suspended culture and biofilm phenotypes including creatine, glutathione, alanine, and choline metabolism and the tricarboxylic acid cycle. These unique metabolite and pathway differences have the potential to serve as targets for *P. aeruginosa* and specifically biofilm diagnosis and biofilm control in synovial fluid.

**Author Summary:** Joint replacement surgery is a common procedure frequently required in later stages of life due to damage in the joint. Over one million joint replacement surgeries are performed annually with rates increasing every year. A devastating complication associated with joint replacement is the development of infection around the implant device in the joint space, known as a periprosthetic joint infection. Bacteria in the joint space can form a biofilm, which is a gel-like matrix encasing the cells that increases resistance to treatment and exacerbates chronic infections. A particular challenge for the diagnosis of biofilm-mediated periprosthetic joint infections is the slowly growing nature of biofilm-mediated phenotypes, resulting in frequent failure to detect these bacteria by clinical microbiological culturing methods. Small molecule metabolites are uniquely produced by strains of bacteria in the biofilm versus planktonic or suspended culture phenotype. Identification of metabolites as specific markers of infection and biofilm could allow a new culture-free diagnostic approach to diagnose infection by biofilm. Furthermore, knowledge of metabolic pathway populations in biofilm in joint fluid could point to specific targets to prevent biofilm formation in the joint space.

## Introduction

Total joint arthroplasty, or joint replacement, is one of the most successful surgical procedures and has been shown to substantially improve patient quality of life (1). Patients often require joint arthroplasty in the later stage of life due to arthritis, joint pain, swelling, lack of blood flow, and trauma. It is a common procedure with over one million hip and knee replacements performed in 2015 (1). Due in part to the increasing rates of arthritis, obesity, and increasing life span, occurrence of joint replacement is expected to increase every year and six-fold by 2040 (2). However, periprosthetic joint infections (PJI) are a leading cause of surgery failure, occurring in 1-2% of cases. PJI pose a major challenge associated with morbidity and mortality in patients and cause a significant economic burden expected to exceed 1.62 billion in the US annually (3–5).

PJI are most commonly caused by *Staphylococci*, but also Gram negative bacteria, such as *P. aeruginosa*, which is of clinical importance due to the difficulty in treating these pathogens (6). The current diagnostic strategy for PJI outlined by the Musculoskeletal Infection Society (MSIS) involves a multi-pronged approach of varied tests on blood, synovial fluid, and tissue samples (3, 7). Diagnosis by traditional approaches is difficult and time-consuming due to collection of multiple specimens, slow growing variants, lengthy culturing methods, and a high rate of false negative cultures (7).

A major factor contributing to the burden of PJI is bacterial biofilm formation where cells can lie dormant, protected by an extracellular matrix against antibiotics and immune responses (3). The presence of the biofilm phenotype contributes to diagnostic and treatment challenges, including considerably increased antibiotic tolerance and treatment failure leading to additional surgeries, amputation, and death is not uncommon (3). Therefore, there is a critical need for the development of new approaches for a more accurate and rapid pathogen- and phenotype-specific diagnosis of PJI and new treatment strategies to prevent or control biofilms in the joint space.

*P. aeruginosa* has been shown to differentially utilize metabolic pathways in the planktonic and biofilm states (8). Therefore, a metabolomics approach has the potential to identify unique metabolites as markers of *P. aeruginosa* infection, and even distinguish between planktonic or suspended culture and biofilm phenotypes. Metabolomics has been applied in select cases to study joint fluid for other conditions such as osteoarthritis and osteochondrosis, but not yet to investigate the potential for *P. aeruginosa* or biofilm specific metabolites (9–12). Because detailed information is lacking about *P. aeruginosa* metabolism and cellular activities in the synovial fluid environment, we aim to evaluate the potential of a metabolomics approach to detect unique metabolites as markers of *P. aeruginosa* PAO1, including *P. aeruginosa* PAO1 as a biofilm, and determine metabolic pathway differences in the suspended culture and biofilm phenotypes in synovial fluid.

Here we utilize 2D NMR spectroscopy for an untargeted metabolomics analysis of bovine synovial fluid (BSF) and *P. aeruginosa* grown in both the suspended culture and lawn biofilm phenotypes in BSF. Our goal is to identify metabolites unique to *P. aeruginosa* cultures compared to the uninoculated BSF control as well as *P. aeruginosa* phenotypic specific metabolites to discriminate between suspended and biofilm cultures. Moreover, metabolite changes between suspended and biofilm states can then be mapped to particular metabolic pathways in search for novel targets for biofilm control.

## Results

### PAO1 growth in 50% BSF/PBS

*P. aeruginosa* wild-type reference strain PAO1 was inoculated in 50% BSF/PBS for planktonic or suspended culture and lawn biofilm growth (13). BSF was used as the sole nutrient source and has been used as a model for bacterial growth in human synovial fluid (14–18). BSF was incubated similarly to the cultures without inoculation as a control. PAO1 culture flasks appeared bright yellow/green and opaque compared to the uninoculated BSF control, potentially correlated with high levels of pyoverdine production (**S1 Fig**) (19). Due to the propensity of both Staphylococci and Gram-negative bacteria to form biofilm-like aggregates in synovial fluid, planktonic cultures are referred to as “suspended cultures” due to the likelihood of aggregate formation in the media (13). Lawn biofilms are known to generate large amounts of biomass and have been used as models of bacterial growth on soft surfaces such as mucosal surfaces and tissue (20, 21). The lawn biofilms grew evenly over the surface of 50% BSF/PBS agar and appeared bright yellow/green compared to the uninoculated BSF control (**S2 Fig**).

After PAO1 growth in BSF, suspended and lawn biofilm cultures yielded similar total cell numbers as determined by CFUs (**S3 Fig**). A total of four independent replicates were included for controls and biofilm cultures and three independent replicates were included for suspended cultures. Due to the appearance of small colony variants in the CFU plates of one suspended culture replicate indicating cell populations of multiple phenotypes, this culture was excluded from analysis. By contrast, CFUs for all other cultures showed a uniform cellular phenotype with colonies of similar color, shape, and size.

### Untargeted metabolomics analysis of BSF and *P. aeruginosa* in suspended culture and biofilm phenotypes

The metabolic differences between the uninoculated BSF control and *P. aeruginosa* PAO1 in the suspended culture and static lawn biofilm phenotypes grown in BSF were identified by untargeted NMR-based metabolomics. For all samples, their 2D ^13^C-^1^H HSQC and 2D ^1^H-^1^H TOCSY spectra were processed and uploaded to the COLMARq web server for metabolite identification and quantification as previously described (8). A representative ^13^C-^1^H HSQC spectrum for each group (control, suspended, and biofilm) is shown as a color-coded overlay in **Fig 1**. The spectra display a large number of distinct cross-peaks belonging to many detectable metabolites in each sample. A sizeable number of HSQC cross-peaks (~50) were unique to each group, which reflects the unambiguous presence of metabolites that are not only unique to *P. aeruginosa* compared to the uninoculated control, but also specific for suspended and biofilm phenotypes.

**Fig 1.**
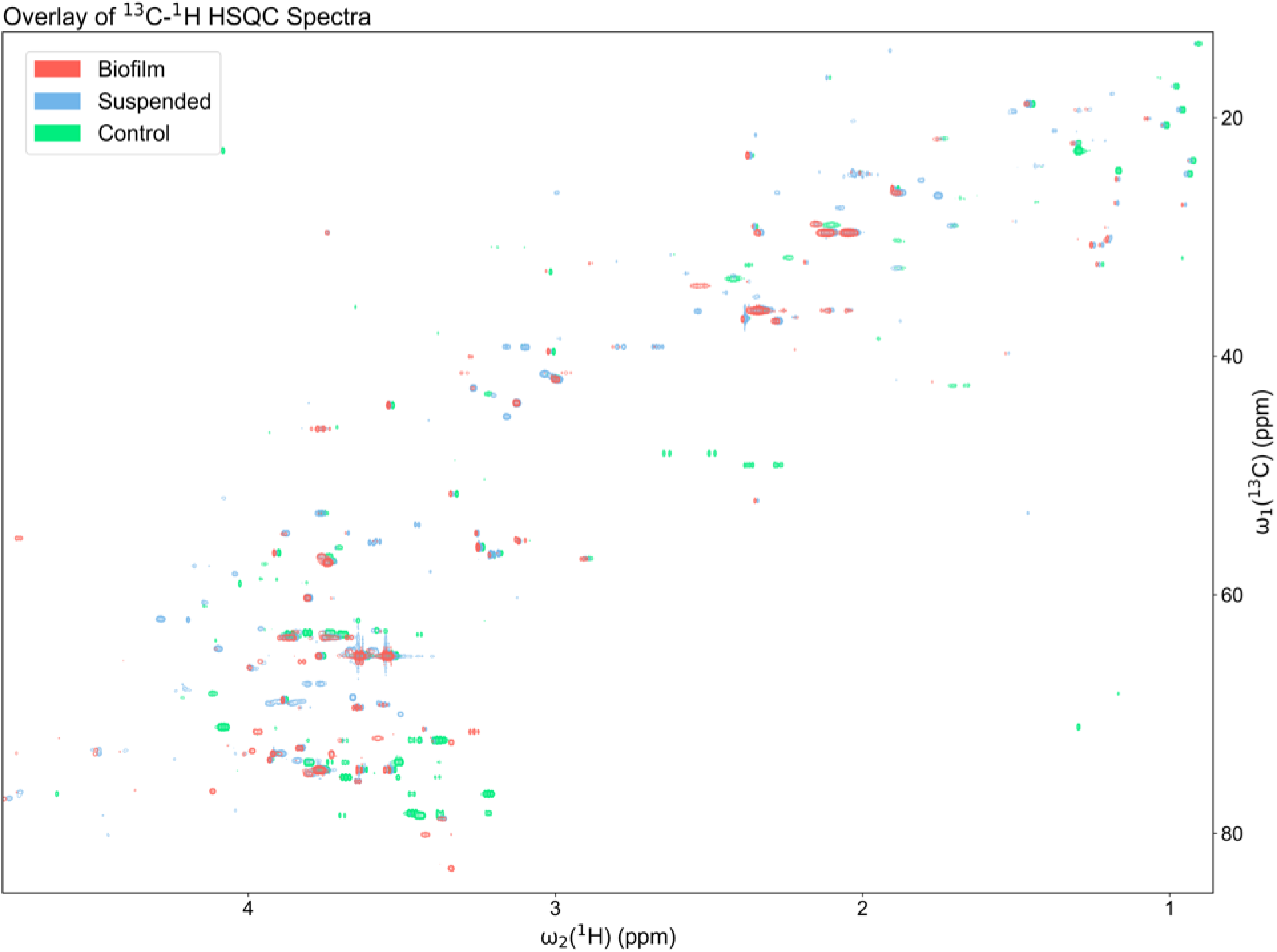
Overlay of a representative region of the 2D ^13^C-^1^H HSQC spectra of a representative uninoculated BSF control (green; bottom), suspended culture (blue; middle), and biofilm (red; top) sample. Samples are all at ^13^C natural abundance and each shifted 0.01 ppm down-field (left) for visualization. Unique peaks were detected between each sample indicating metabolic differences between the BSF control and *P. aeruginosa* cultures grown in BSF as suspended culture and biofilm phenotypes.

### Multivariate analysis of metabolomics data by PCA

Quantitative changes in metabolites between the three groups were evaluated using principal component analysis (PCA) as an unsupervised multivariate statistical analysis approach (**Fig 2**). The samples clustered tightly within their groups indicating high reproducibility. The groups were also well separated with no overlap of the 95% confidence regions indicating distinct metabolite differences between the groups. The PC1 comprised 68.4% of the variance in the data, mostly corresponding to separation between the *P. aeruginosa* cultures and BSF control. The PC2 comprised 21.5% of the variance in the data dominated by the mean separation between the suspended and lawn biofilm cultures. With just three to four samples per cohort, the spread within each cohort in the PCA score plot is notably small indicating high overall reproducibility with good, unsupervised inter-cohort and phenotype discrimination.

**Fig 2.**
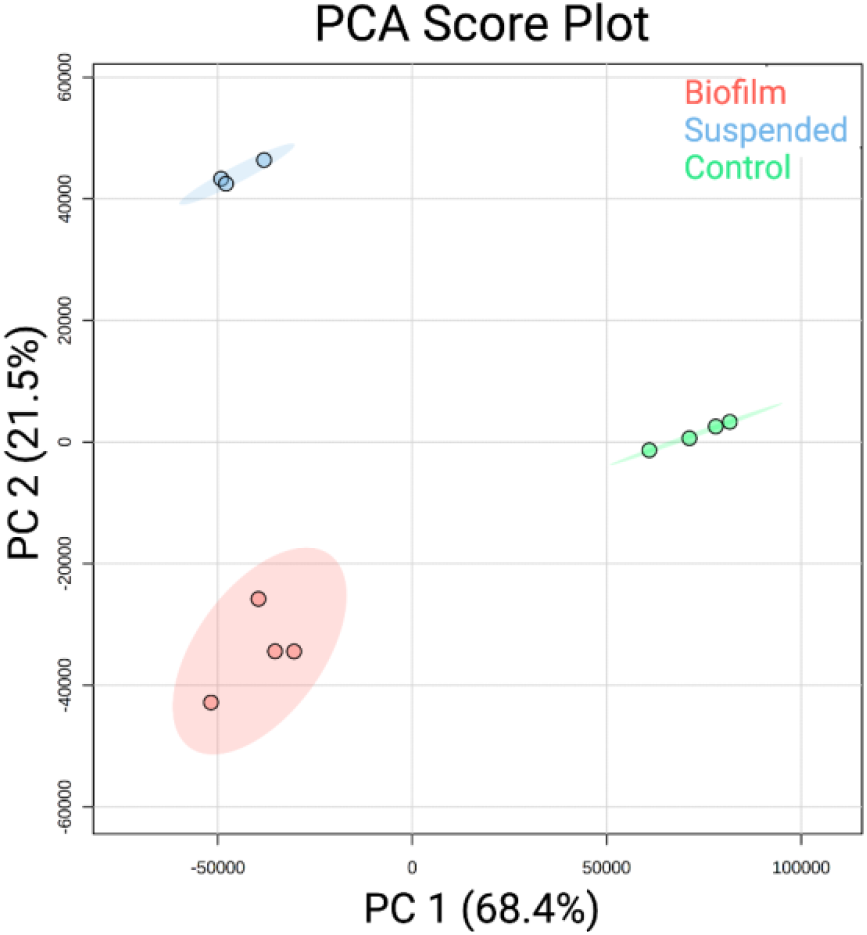
The two-dimensional score plot for principal component analysis (PCA) of the uninoculated BSF controls (green) (n=4), suspended cultures (blue) (n=3), and lawn biofilms (red) (n=4). The PCA is based on the quantitation of identified metabolites showing clustering of sample cohorts with no overlap of the ellipses (ellipses represent 95% confidence intervals), displaying good separation between and repeatability within cohorts of samples.

### Metabolites uniquely detected in the uninoculated BSF control

A total of 51 different metabolites were detected across all samples. Of these, 30 metabolites were identified in the uninoculated BSF control. This includes 12 metabolites that were identified only in the control comprising several amino acids, lactic acid, and glucose (**Fig 3**).

**Fig 3.**
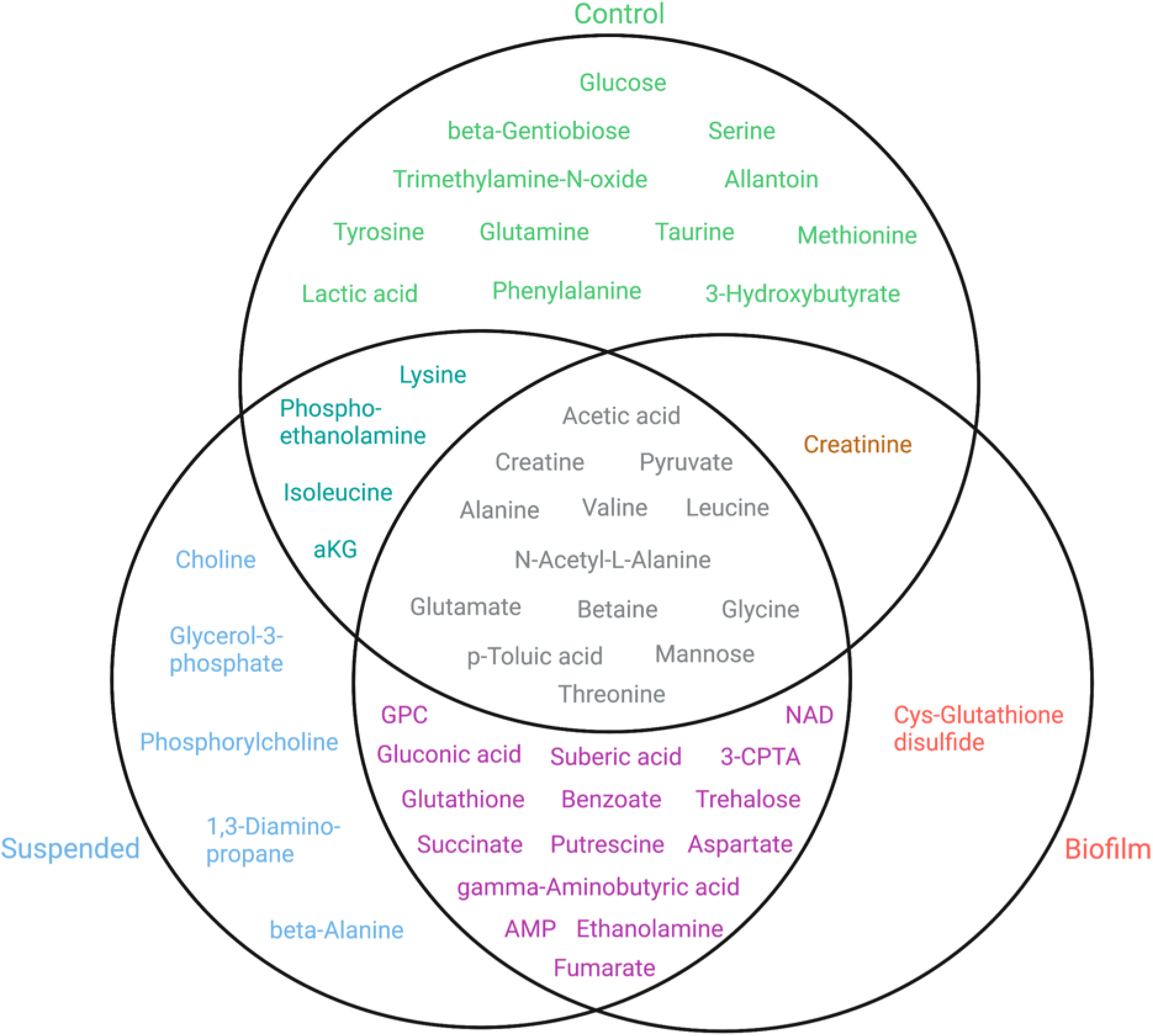
A Venn diagram of metabolites identified in the uninoculated BSF control (green), suspended culture (blue), and lawn biofilm (red) samples. The overlap of circles indicates metabolites detected in multiple sample types (aKG = alpha-ketoglutarate, GPC = glycerophosphocholine, NAD = nicotinamide adenine dinucleotide, AMP = adenosine monophosphate, 3-CPTA = 3-carboxypropyl-trimethyl-ammonium, Cys = cysteine).

### Metabolites uniquely detected with *P. aeruginosa* in both growth phenotypes

A total of 39 metabolites were identified in the suspended culture and lawn biofilm *P. aeruginosa* samples. The relative metabolite quantities are visualized by means of a heatmap in **Fig 4**. Hierarchical clustering by Ward’s method and Euclidean distance show biofilm and suspended culture samples were distinctly clustered into two groups. A total of 21 metabolites were identified to be present in the *P. aeruginosa* cultures that were absent in the uninoculated BSF control (**Fig 3**).

**Fig 4.**
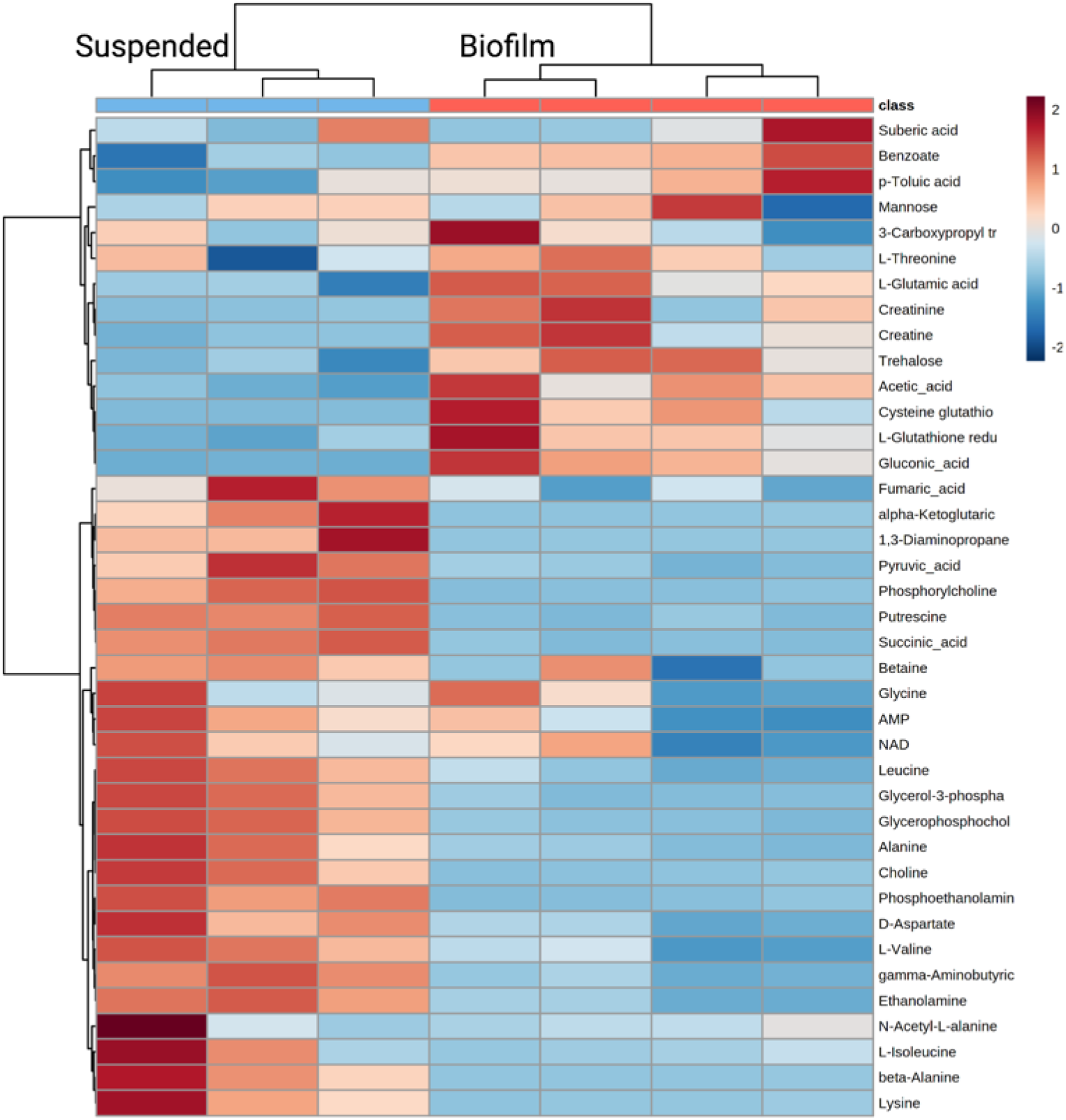
A quantitative visualization of metabolites identified in *P. aeruginosa* suspended culture and lawn biofilm samples by a heatmap. The map uses hierarchical clustering to accurately cluster samples into their respective cohorts (see text). The color scale shows metabolite fold changes between cohorts.

Of these metabolites, 15 were identified in both the suspended and biofilm cultures as non-phenotype specific indicators of *P. aeruginosa* in bovine synovial fluid (**Fig 3**). While present in both phenotypes, some of these metabolites showed large differences in their abundance between the suspended culture and biofilm phenotypes. Metabolites that are significantly higher in suspended culture include putrescine (fold change (FC) biofilm/suspended = 0.04, *p* = 3.12 × 10^-6^), succinate (FC = 0.08, *p* = 6.25 × 10^-6^), glycerophosphocholine (FC = 0.13, *p* = 2.88 × 10^-4^), gamma-aminobutyric acid (FC = 0.31, *p* = 9.96 × 10^-5^), aspartate (FC = 0.31, *p* = 1.96 × 10^-3^), fumarate (FC = 0.32, *p* = 2.87 × 10^-2^), and ethanolamine (FC = 0.39, *p* = 1.76 × 10^-4^) (**Fig 4**, **S1 Table**). Several of these metabolites, namely gamma-aminobutyric acid, fumarate, and ethanolamine also have isolated unique peaks in the 1D ^1^H NMR spectra (**Fig 5a**). Metabolites that are significantly higher in biofilm include gluconic acid (FC = 4.90, *p* = 7.08 × 10^-3^), glutathione (FC = 3.53, *p* = 2.66 × 10^-2^), and trehalose (FC = 3.18, *p* = 9.20 × 10^-3^) (**Fig 4**, **S1 Table**).

**Fig 5.**
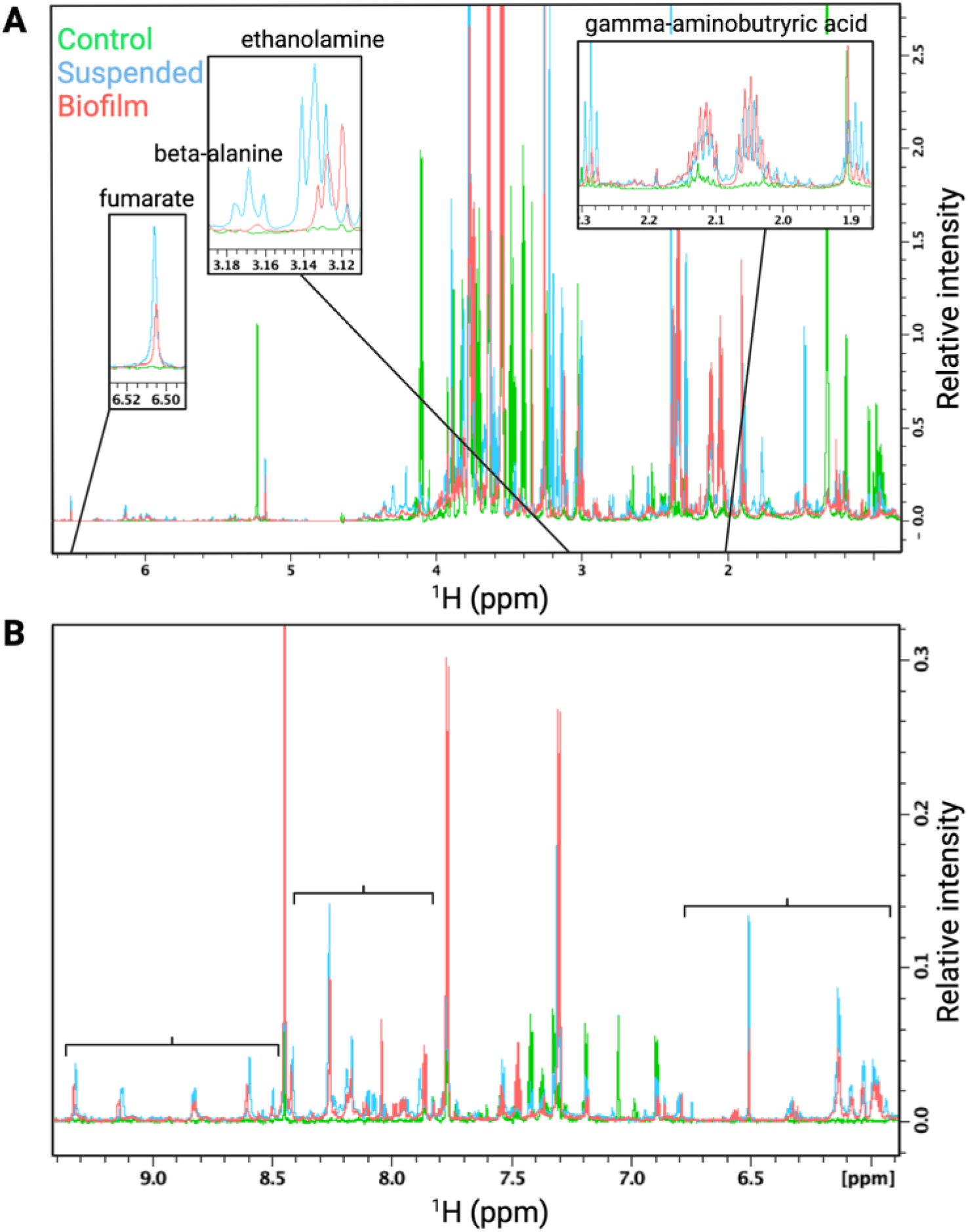
Overlay of regions of a representative 1D ^1^H NMR spectrum from the uninoculated BSF control (green), suspended culture (blue), and lawn biofilm (red) samples. Spectrum (**A**) shows zoomed in regions for metabolites with unique peaks in the 1D spectrum. Fumarate, ethanolamine, and gamma-aminobutyric acid are uniquely present in *P. aeruginosa* suspended and lawn biofilm cultures and absent in the uninoculated control. Beta-alanine was uniquely present in the 1D in the suspended culture. The brackets in spectrum (**B**) indicate three spectral regions containing NMR signals unique to the *P. aeruginosa* suspended and lawn biofilm cultures and void of signals from the uninoculated BSF control.

### Metabolites uniquely detected with *P. aeruginosa* in either the suspended culture or biofilm phenotype

A total of six metabolites, whose relative abundance is depicted as box plots in **Fig 6**, were also exclusively detected in only *P. aeruginosa* suspended culture or lawn biofilms. Therefore, these metabolites unambiguously indicate the presence of *P. aeruginosa* suspended culture or biofilm phenotypes in synovial fluid and have potential as phenotype-specific markers. Metabolites uniquely detected in the *P. aeruginosa* suspended culture phenotype include phosphorylcholine, choline, 1,3-diaminopropane, glycerol-3-phosphate, and beta-alanine. Among them, beta-alanine also has a well-isolated peak in the 1D ^1^H NMR spectra and, hence, could be monitored in such samples solely based on 1D ^1^H NMR. The one metabolite exclusively detected in the *P. aeruginosa* biofilm phenotype is cysteine-glutathione disulfide (**Figs 3** and **7**). Cysteine-glutathione disulfide is a fair metabolite match in the spectra with 3 HSQC peaks that are uniquely matched to this metabolite.

**Fig 6.**
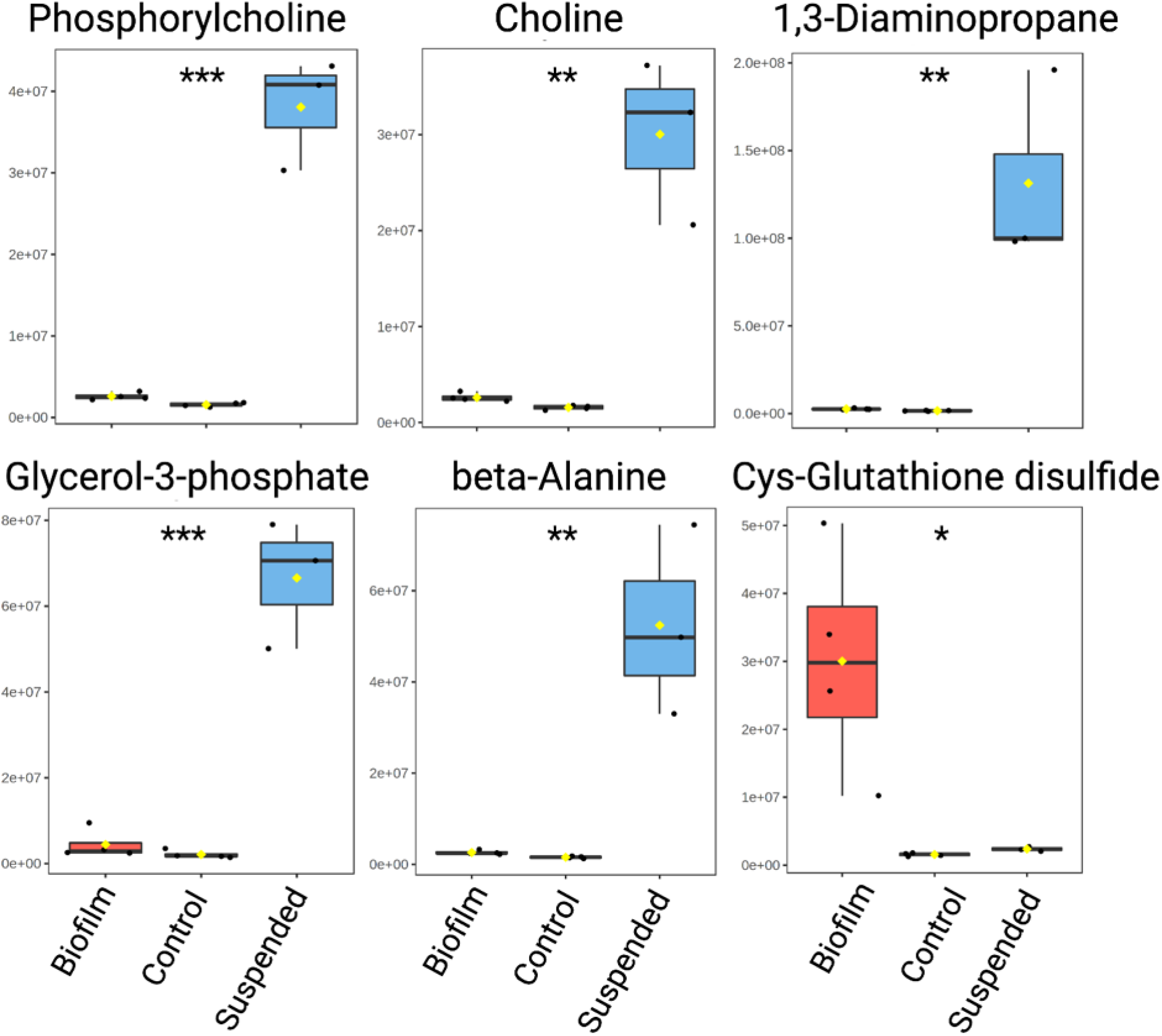
Uniquely detected metabolites in suspended or lawn biofilm cultures. The box plots represent a metabolite quantity analysis between the uninoculated BSF control (green) (n=4), suspended (blue) (n=3), and lawn biofilm (red) (n=4) cultures. The black circles represent independent sample values, boxes represent upper and lower quartiles, black bars represent median, yellow diamonds 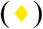 represent the mean value, and whiskers represent minimum and maximum values, and asterisks denote significance between suspended culture and biofilm sample cohorts (* *p* 0.05, \*\**p*<0.01, \*\*\**p*<0.001 by unpaired, two-tailed *t*-test). (Cys = cysteine)

**Fig 7.**
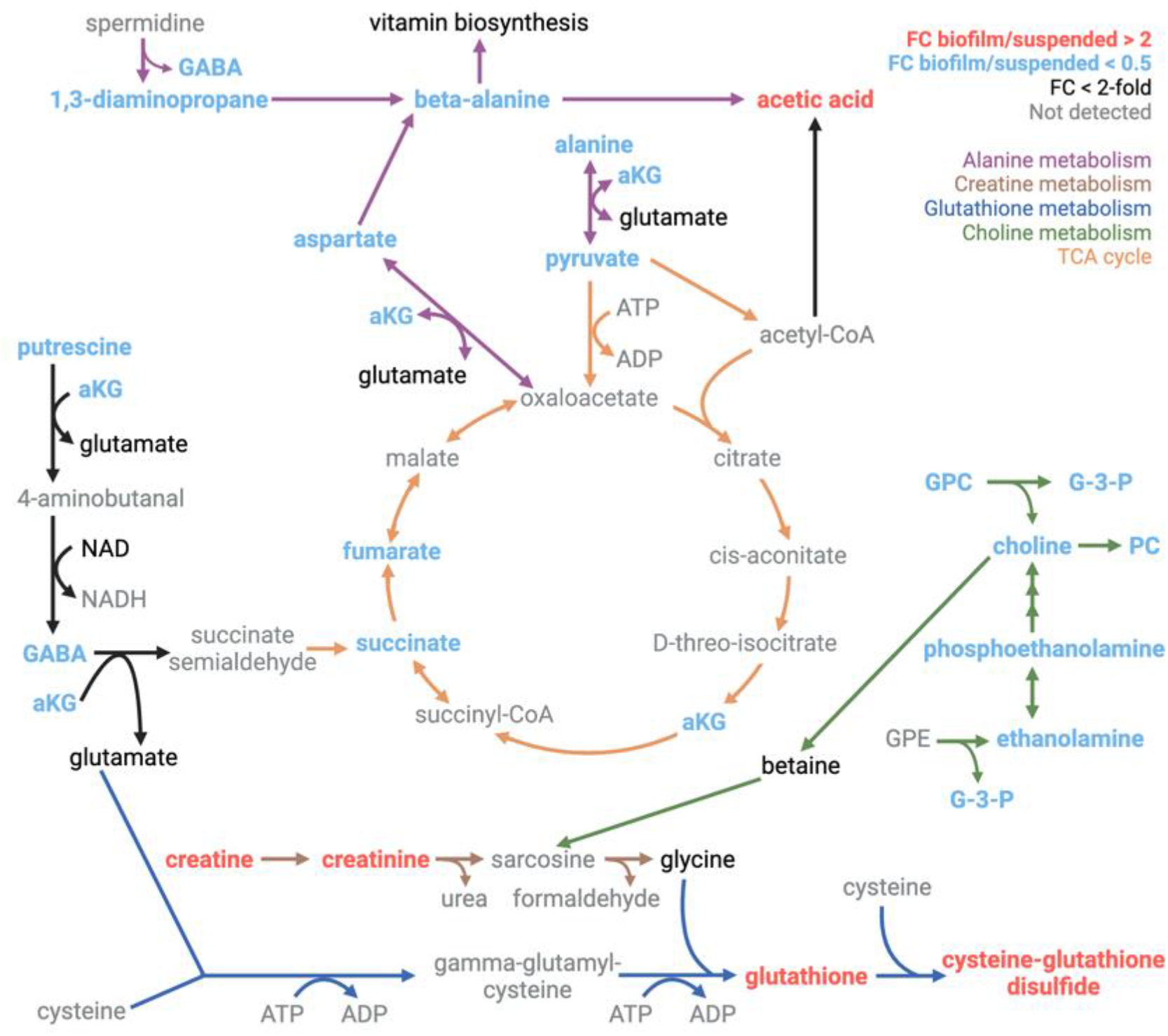
Metabolic pathway map displaying significant metabolite quantity differences between the suspended and lawn biofilm cultures. Metabolites with a fold change (FC) (biofilm/suspended) > 2 are red indicating higher levels in biofilm and FC < 0.5 are blue indicating higher levels in suspended culture. Pathways showing significant changes between suspended culture and biofilm have the arrows color-coded including alanine metabolism (purple), creatine metabolism (brown), glutathione metabolism (dark blue), choline metabolism (green), and the TCA cycle (orange). Metabolites that were detected but the fold change is less than two-fold are shown in black and metabolites not detected are shown in gray (GABA = gamma-aminobutyric acid, aKG = alpha-ketoglutarate, ATP = adenosine triphosphate, ADP = adenosine diphosphate, NAD = nicotinamide adenine dinucleotide, NADH = nicotinamide adenine dinucleotide reduced, GPC = glycerophosphocholine, G-3-P = glycerol-3-phosphate, PC = phosphorylcholine, GPE = glycerophosphoethanolamine).

### NMR spectral peaks of unknown metabolites unique to *P. aeruginosa* and each phenotype

In addition to the unique metabolites discussed above, we observed many 2D as well as 1D NMR spectral peaks belonging to unknown metabolites. 54 unknown peaks were unique to *P. aeruginosa* compared to the uninoculated BSF control and 25 were unique to the suspended culture or biofilm phenotypes (**Table 1**). Some peaks could be uniquely detected in 2D but not 1D NMR spectra because of high peak overlap in 1D and other peaks could only be detected in the 1D due to its higher sensitivity compared to 2D.

**Table 1.** Peaks uniquely detected in the 2D ^13^C-^1^H HSQC and 1D ^1^H NMR spectra in *P. aeruginosa* suspended and lawn biofilm cultures but absent in the uninoculated BSF control. The ‘*’ indicates peaks that are uniquely detected in both the 2D and 1D spectra.

| Unique peaks in *P. aeruginosa* (suspended & biofilm) (ppm) |  |  |  | Unique peaks in suspended (ppm) |  |  |  | Unique peaks in biofilm (ppm) |  |
| --- | --- | --- | --- | --- | --- | --- | --- | --- | --- |
| $^1\text{H}$ | $^{13}\text{C}$ | | $^1\text{H}$ | $^1\text{H}$ | $^{13}\text{C}$ | | | $^1\text{H}$ | $^{13}\text{C}$ |
| 1.21 | 29.19 |  | 1.08 | *1.82 | 25.25 |  |  | 1.30 | 30.18 |
| 1.26 | 30.69 |  | 2.39 | *3.17 | 45.07 |  |  | 1.56 | 25.15 |
| 1.27 | 19.32 |  | 5.98 | 3.42 | 58.06 |  |  | 2.96 | 41.41 |
| *1.51 | 28.69 |  | 6.08 | 3.46 | 54.11 |  |  | 3.26 | 71.48 |
| *1.53 | 39.76 |  | 6.13 | 3.61 | 55.67 |  |  | 3.28 | 40.02 |
| *2.05 | 36.17 |  | 7.30 | 3.67 | 68.59 |  |  | *3.34 | 72.36 |
| 2.12 | 36.16 |  | 7.77 | 3.82 | 67.48 |  |  | 3.34 | 80.09 |
| *2.34 | 29.66 |  | 8.04 | *4.20 | 62.10 |  |  | 3.34 | 82.93 |
| 3.26 | 54.83 |  | 8.26 | 4.25 | 73.78 |  |  | 3.42 | 80.11 |
| 3.27 | 42.67 |  |  |  |  |  |  | 3.45 | 58.84 |
| 3.56 | 69.26 |  |  |  |  |  |  | 3.53 | 71.70 |
| 3.68 | 54.82 |  |  |  |  |  |  | 3.58 | 72.02 |
| 3.73 | 60.32 |  |  |  |  |  |  | 3.97 | 71.47 |
| 3.75 | 29.65 |  |  |  |  |  |  | 3.98 | 73.08 |
| 3.80 | 63.61 |  |  |  |  |  |  | 4.45 | 81.19 |
| 4.11 | 64.52 |  |  |  |  |  |  | 4.62 | 72.02 |
| 4.37 | 86.45 |  |  |  |  |  |  |  |  |
| 4.82 | 84.36 |  |  |  |  |  |  |  |  |
| 5.24 | 103.96 |  |  |  |  |  |  |  |  |
| *8.17 | 155.41 |  |  |  |  |  |  |  |  |

Altogether, 20 2D HSQC peaks were present in *P. aeruginosa* cultures and absent in the control. Of these, 5 were isolated in the 1D ^1^H spectra and there were an additional 9 unique peaks present in the 1D spectra. There were 9 2D HSQC peaks unique to the suspended culture phenotype, with 3 of these also isolated in the 1D spectra. There were 16 2D HSQC peaks unique to the biofilm phenotype, but only 1 of them also isolated in the 1D spectra (**Table 1**). In addition, entire spectral regions of the 1D ^1^H spectra have peaks that were exclusively detected in *P. aeruginosa* cultures but absent in the BSF control. These regions are down-field shifted and they cover ranges 5.94-6.87, 7.89-8.43, and 8.48-9.34 ppm (**Fig 5b**). Metabolites with resonances in these regions typically contain aromatic rings, carbon double bonds, or aldehyde groups.

### Metabolites and pathways differing significantly between suspended culture and biofilm

Mapping the quantitative metabolite differences between *P. aeruginosa* suspended and biofilm lawn cultures to metabolic pathways could provide insight into pathways that are differentially utilized in each growth mode in synovial fluid (**Fig 7**). Metabolites found to be significantly increased in the biofilm compared to suspended culture phenotypes include cysteine-glutathione disulfide (FC = 12.64, *p* = 3.83 × 10^-2^) and glutathione (FC = 3.53, *p* = 2.66 × 10^-2^) which are involved in glutathione metabolism. Creatine (FC = 3.43, *p* = 4.80 × 10^-2^) and creatinine (FC = 4.80, *p* = 6.89 × 10^-2^) were also found with increased abundance in the biofilm lawn. Although creatinine showed a FC = 4.80, the *p*-value was not significant *(p* = 0.07). Additional replicates are needed to more robustly assess if this metabolite difference is statistically significant. Gluconic acid (FC = 4.90, *p* = 7.08 × 10^-3^) and trehalose (FC = 3.18, *p* = 9.20 × 10^-3^) are carbohydrate-related metabolites that were found to be significantly increased in the biofilm compared to suspended culture.

Many metabolites were found to be statistically significantly decreased in the biofilm lawn compared to the suspended cultures (*p*<0.05), including metabolites belonging to pathways of choline metabolism, alanine metabolism, and the TCA cycle (**Fig 7**). In particular, metabolites related to choline metabolism including phosphorylcholine (FC = 0.07, *p* = 1.22 × 10^-4^), glycerol-3-phosphate (FC = 0.07, *p* = 4.13 × 10^-4^), choline (FC = 0.09, *p* = 1.15 × 10^-3^), phosphoethanolamine (FC = 0.11, *p* = 4.08 × 10^-5^), glycerophosphocholine (FC = 0.13, *p* = 2.88 × 10^-4^), and ethanolamine (FC = 0.39, *p* = 1.76 × 10^-4^) were significantly decreased in biofilm. Metabolites associated with alanine metabolism that were significantly decreased in biofilm were 1,3-diaminopropane (FC = 0.02, *p* = 4.99 × 10^-3^), putrescine (FC = 0.04, *p* = 3.12 × 10^-6^), beta-alanine (FC = 0.05, *p* = 4.36 × 10^-3^), alanine (FC = 0.16, *p* = 3.31 × 10^-3^), gamma-aminobutyric acid (FC = 0.31, *p* = 9.96 × 10^-5^), and aspartate (FC = 0.31, *p* = 1.96 × 10^-3^). Intermediates of the TCA cycle were also significantly decreased in biofilm lawns compared to suspended cultures, including alpha-ketoglutarate (FC = 0.07, *p* = 3.19 × 10^-3^), succinate (FC = 0.08, *p* = 6.25 × 10^-6^), pyruvate (FC = 0.22, *p* = 1.88 × 10^-3^), and fumarate (FC = 0.32, *p* = 2.87 × 10^-2^).

## Discussion

While metabolomics has been recognized as a potentially useful diagnostic approach for PJI, it has not yet been fully evaluated (22). Here we find that NMR-based monitoring of metabolite markers has strong promise for further development as a method of detection of *P. aeruginosa* suspended culture and biofilm phenotypes in synovial fluid. A total of 21 metabolites, 54 unknown metabolite peaks, and distinct 1D ^1^H NMR spectral regions were uniquely detected in *P. aeruginosa* in both phenotypes compared to BSF alone. Five metabolites were uniquely detected in suspended culture whereas only cysteine-glutathione disulfide was uniquely detected in biofilm (**Fig 6**). Cysteine-glutathione disulfide, with its potential as a marker for the presence of *P. aeruginosa* biofilm, is formed upon oxidative stress of glutathione, but little is known about its specific role in *P. aeruginosa* metabolism (23, 24). It has been detected in the saliva and plasma of humans and has been shown to play a role in preventing toxicity of certain drugs in mammalian cells (25–27).

The 54 unknown metabolite peaks (**Table 1**), even without knowing the metabolites they belong to, could also be used as potential indicators of suspended culture or biofilm by 2D or 1D NMR. In fact, future research to identify these unknown metabolites should lead to the identification of additional metabolite markers for *P. aeruginosa* in the suspended culture or biofilm phenotypes along with new insights on changes in metabolic pathways associated with each phenotype.

Utilizing the unique, but less detailed 1D ^1^H NMR spectral regions (**Fig 5b**) comprising many different peaks could serve as a robust diagnostic strategy in practice. Spectral regions can be observed by more cost-effective low-field NMR methods that lack the resolution of NMR at high magnetic field (28). Therefore, these spectral regions have the potential as rapid markers for identification of *P. aeruginosa* in synovial fluid or joint aspirates. If these potential diagnostic markers are transferrable to *ex vivo* clinical PJI joint fluid aspirates, these metabolites and peaks could become powerful markers for the detection of *P. aeruginosa*, and biofilm specifically, in synovial fluid to complement traditional culturing methods that are time consuming (days) and have a high rate of false negatives (7).

Due to its reproducibility, quantitative capabilities, and ability to detect a large number of metabolites simultaneously, NMR is well-suited for untargeted metabolomics analyses (29, 30). Our analysis shows that 2D NMR-based metabolomics provides a higher level of detail than 1D ^1^H NMR allowing the confident identification of many more metabolites along with 36 additional metabolite cross-peaks belonging to unknown metabolites that are uniquely detected in *P. aeruginosa* cultures compared to the uninoculated control. However, the routine use of 2D NMR in clinical diagnostics is hindered by prolonged measurement times on the order of hours. On the other hand, 1D NMR spectra can be rapidly collected within minutes. While use of NMR is still relatively limited in clinical diagnostics, there are increasing numbers of hospitals with NMR machines for metabolomics analysis of patient samples, for example for cardiovascular disease, diabetes, and cancer (31). These newly identified metabolites may also be translated to other more targeted methods that are faster and less costly than NMR, such as biosensors or immunoassays (32, 33).

The unique metabolite markers were identified from analysis of cohorts of three and four independent suspended culture and biofilm samples, respectively. Although the total number of replicates used in this study is small, the samples have high reproducibility and distinct inter-cohort and phenotype separation as manifested in distinct clustering in the PCA score plot (**Fig 2**). While we found many metabolites unique to each culture and the control, it is possible that they may be weakly present on the opposing cohorts, but below the NMR limit of detection. Further studies with increased sample size and efforts to identify the metabolite markers by a more sensitive complementary method such as mass spectrometry can be used to further corroborate these findings.

Our results provide a proof of principle for detection of unique *P. aeruginosa* suspended culture and biofilm metabolites in synovial fluid. While BSF is used as a model of human synovial fluid for bacterial growth (14–18), it is naturally devoid of complicating factors such as an active immune response that can significantly modify the local environment. Measurements of human samples to determine the metabolite background of human synovial fluid and *P. aeruginosa* infection *ex vivo* are needed to further evaluate the diagnostic utility of this method. Since many species of bacteria have been shown to preferentially utilize different metabolic pathways (34), different species such as *Staphylococci* may produce distinct metabolites in this environment and thus NMR has the potential to identify species specific metabolites. It is also important to test our approach on *P. aeruginosa* clinical strains and other pathogens to determine if the metabolite markers are unique to *P. aeruginosa* specifically or more widely indicative of the presence of bacteria.

We also report significant differences in several metabolic pathways between *P. aeruginosa* suspended culture and biofilm phenotypes when grown in BSF (**Fig 7**). Metabolic pathways significantly increased in biofilm include creatine and glutathione metabolism. Creatinine has been shown to inhibit replication in bacteria including *P. aeruginosa* (35) and creatine metabolism is linked to glutathione metabolism through production of glycine (36). Glutathione metabolism in *P. aeruginosa* is known to affect quorum sensing (QS), virulence factor production and secretion, response to oxidative stress, motility, and biofilm formation (37–39). Glutathione has been associated with virulence in *P. aeruginosa* by acting as a redox signal to upregulate the type III secretion system to release effector proteins into host cells (39). Deletion of glutathione biosynthesis genes in PAO1 in two separate studies have linked glutathione to the increased production of virulence factors like pyocyanin and persister cell formation, however they reported conflicting results about glutathione’s effect on increasing or reducing biofilm formation (38, 39). We find glutathione (and related metabolites) in significantly higher concentration in biofilm suggesting it may play a role in the biofilm phenotype. Associated activated pathways in biofilm may reveal new potential enzyme and metabolite targets for regulating biofilm formation.

Several pathways were significantly decreased in biofilm including alanine metabolism, choline metabolism, and the TCA cycle. Alanine metabolism is linked to supplement pantothenate, or vitamin B5, biosynthesis (40) and peptidoglycan for the synthesis of cell walls (41). Choline metabolism is linked to the production of phospholipids for cell membrane synthesis (42). These processes, along with the TCA cycle are important in cellular proliferation. Taken together, the reduced levels of metabolites associated with cellular proliferation and energy metabolism, align with previous reports to indicate a more dormant state of cells in a biofilm. Our findings support that a more metabolically dormant state is relevant for *P. aeruginosa* biofilm grown in BSF (22).

Further investigation of the role of these major pathway changes in biofilm formation are needed. As previously demonstrated, a strategy of metabolic regulation of biofilm formation may be possible by the external supplementation of metabolites that are significantly decreased in biofilm and that are involved in major pathway differences (8). Further investigation is needed to explore these metabolic pathways and related enzymes as potential targets to prevent or reduce biofilm formation in synovial fluid.

As the first study, to our knowledge, of *P. aeruginosa* metabolism grown solely on BSF, we also compared our results to our previous findings of *P. aeruginosa* planktonic and biofilm cultures grown in LB, a rich microbiological media (8). In the uninoculated BSF control we detected a wide range of carbon sources that bacterial pathogens can preferentially catabolize, such as glucose, pyruvate, and 13 amino acids (**Fig 3**) (43). However, both phenotypes exhibit lower growth in BSF, as BSF cultures required triple the liquid volumes and number of agar plates to produce cultures of similar cells numbers to those grown in LB (**S3 Fig**) (8) and fewer total metabolites were detected. The major metabolic pathway differences between planktonic/suspended culture and biofilm phenotypes varied when grown on BSF compared to LB. **S4 Fig** shows a PCA score plot for NMR metabolomics data from cultures grown as suspended cultures and lawn biofilms in both LB and BSF. The plot shows distinct clustering of cultures into their respective cohorts, indicating metabolic differences between each growth mode. There is clear separation between suspended cultures and lawn biofilms regardless of media, but there is a much greater separation between PAO1 cultures grown in LB versus BSF. Variation in metabolic regulation in LB versus BSF reflects the highly adaptable nature of PAO1 metabolism with respect to their nutritional environment and suggests that metabolomics results obtained in different growth media are not necessarily transferrable. It is therefore important to conduct experiments for the development of *in vivo* diagnostics under relevant nutrient conditions, such as using synovial fluid for a more accurate representation of the PJI environment. Nevertheless, in both LB and BSF conditions gluconic acid and trehalose were found in significantly higher concentrations in lawn biofilms compared to suspended cultures. These metabolites may play a role in composing the biofilm matrix as carbohydrates make up a large component of the extracellular matrix (44). High levels of these carbohydrate-related metabolites, therefore, have the potential to uncover the presence of *P. aeruginosa* biofilm under multiple growth conditions.

In conclusion, we identified and fully quantified a wealth of significant metabolite differences between BSF and *P. aeruginosa* in the suspended culture and biofilm phenotypes grown in BSF. This study provides a proof of principle for the use of NMR for identifying unique metabolic markers of *P. aeruginosa* and specifically biofilm in BSF. The observed differences in metabolite content and concentrations can serve the development of diagnostic markers or even therapeutic targets for controlling or preventing biofilms in PJI.

## Material and Methods

### Bacterial strains, growth media, and culturing methods

*P. aeruginosa* strain PAO1 burn wound isolate (45) cultures were grown in lysogeny broth (LB) (Sigma Aldrich) shaking at 220 rpm at 37°C for 24 hours (hrs) to OD_600_ ≈ 1.0. Cultures were diluted in phosphate buffered saline (PBS) to OD_600_ = 0.1 then grown in suspended culture or plated for metabolomics experiments. BSF (Lampire Biological Laboratories) was thawed and spun down 4,300 × *g* for 30min at 25°C prior to use for culturing. PAO1 was grown in suspended culture in 150 mL 50% BSF/PBS broth at 220 rpm at 37°C for 24 hrs (n = 3) and as a biofilm lawn on three 50% BSF/PBS plates (28.4 cm^2^) per sample containing 1.5% (w/v) agar, statically, at 37°C in 5% CO_2_ for 48 hrs (n = 4). No nutrient supplement was used. Control samples without inoculation containing 50% BSF/PBS broth were incubated at 220 rpm at 37°C for 24 hrs (n = 4). Colony-forming units (CFU)/mL (n = 3) and CFU/mL × cm^2^ (n = 3) were measured for suspended and biofilm cultures, respectively, for metabolomics measurements by the microdilution plating technique (46).

### Metabolomics sample preparation

Suspended cultures were harvested by transferring 25 mL aliquots into six conical tubes per sample and centrifugation at 4,300 × *g* for 20 min at 4°C. The pellet was washed by 5 mL PBS, centrifuged again, and transferred into a microcentrifuge tube (Eppendorf). Biofilm cultures were harvested by scraping with a sterile loop and transferring the biomass into two microcentrifuge tubes per plate due to the limited tube capacity. Samples were immediately resuspended in 600 μL cold 1:1 methanol (Fisher) / double distilled H_2_O (ddH_2_O) for quenching. 300 μL of 1.4 mm stainless-steel beads (SSB14B) were added, and cells were homogenized and lysed by a Bullet Blender (24 Gold BB24-AU by Next Advance) at a speed of 8 for 12 min at 4°C (47). An additional 500 μL 1:1 methanol / ddH_2_O was added and the sample was centrifuged at 14,000 × *g* for 10 min at 4°C to remove beads and solid debris. The supernatant was transferred to a 50 mL conical tube and 1:1:1 methanol / ddH_2_O / chloroform (Fisher) was added for a total volume of 24 mL (48, 49). For the BSF control, 1 mL of 50% BSF/PBS without inoculation was transferred to a 50 mL conical tube and similarly 1:1:1 methanol / ddH_2_O / chloroform (Fisher) was added for a total volume of 24 mL. The sample was vortexed and centrifuged at 4,300 × *g* for 20 min at 4°C for phase separation. The aqueous phase was collected and combined for each culture replicate and the methanol content was reduced using rotary evaporation. A centrifugal device with a 3K filter (Pall Microsep) was washed with ddH_2_O three times and the sample was filtered, then frozen and lyophilized overnight. For NMR measurements, the samples were re-suspended in 200 μL NMR buffer (50 mM sodium phosphate buffer in D_2_O at pH 7.2 with 0.1 mM DSS (4,4-dimethyl-4-silapentane-1-sulfonic acid) for referencing). A centrifugal device with a 3K filter (Pall Nanosep) was washed three times with D_2_O and the NMR sample was filtered at 14,000 rpm for 30 min at 4°C. An additional 50uL of NMR buffer was added to the filter, centrifuged again to wash, and combined for a total 250 μL of sample that was transferred to a 3 mm NMR tube with a Teflon cap and sealed with parafilm.

### NMR experiments and processing

NMR spectra were collected at 298 K on a Bruker AVANCE III HD 850 MHz solution-state spectrometer equipped with a cryogenically cooled TCI probe. 1D ^1^H spectra (Bruker pulse program “zgesgppe”) were collected with 16,384 complex points for a measurement time of around 4 min. The spectral width was 13587.0 Hz, and the number of scans was 32. 2D ^1^H-^1^H TOCSY spectra were collected (Bruker pulse program “dipsi2ggpphpr”) with 256 complex t1 and 2048 complex t2 points for a measurement time of 4 hrs. The spectral widths along the indirect and direct dimensions were 10,202.0 and 10,204.1 Hz and the number of scans per t1 increment was 14. 2D ^13^C-^1^H HSQC spectra (Bruker pulse program “hsqcetgpsisp2.2”) were collected with 512 complex t1 and 2048 complex t2 points for a measurement time of 16 hrs. The spectral widths along the indirect and direct dimensions were 34206.2 and 9375.0 Hz and the number of scans per t1 increment was 32. The transmitter frequency offset values were 75 ppm in the ^13^C dimension and 4.7 ppm in the ^1^H dimension for all experiments. NMR data was zero-filled four-fold in both dimensions, apodized using a cosine squared window function, Fourier-transformed, and phase-corrected using NMRPipe (50).

### NMR-based metabolomics data analysis

2D HSQC and TOCSY spectra were uploaded to the new COLMARq web server (http://spin.ccic.osu.edu/index.php/quan/index) for peak picking, peak alignment, metabolite identification, metabolite quantification via Gaussian peak fitting, spectral normalization via global scaling based on the average, median 30% peak volume ratios between an arbitrarily selected reference spectrum, and univariate statistical analysis between cohorts (51, 52). Multivariate statistical analysis, hierarchical clustering analysis and heatmap visualization, and metabolite box plot analysis was performed via MetaboAnalyst (53). Metabolites were mapped to pathways via the KEGG PATHWAY database (54) and MetaCyc (24).

### Statistical analysis

All assays were performed in at least three independent replicates. Two-tailed unpaired Student’s t-tests were used for significant differences between groups. *P*-values below 0.05 were considered statistically significant. Metabolomics results were checked for multiple comparisons testing using the Benjamini-Hochberg false discovery rate (FDR) test (55). All error bars represent one standard deviation.

## Acknowledgments

This work was supported by the National Institutes of Health (grants R01GM124436 (to P.S.) and R35GM139482 (to R.B.)) and by the Pilot Program Funding from the Department of Microbial Infection and Immunity in the College of Medicine at OSU. All NMR experiments were performed at the CCIC NMR facility at The Ohio State University.

## Author Contributions

Conceptualization and design: AL, LB, AS, RB, and PS. Investigation: AL. Data analysis: AL and DL. Figures: AL. Writing of the original draft: AL, RB, and PS. All authors read and approved the final version of the manuscript.

## Supporting information

**S1 Fig. Photos of shaking cultures after 24 hrs of incubation show the 50% BSF/PBS culture inoculated with PAO1 (A) and uninoculated as a control (B).** The PAO1 culture is bright yellow/green and opaque.

**S2 Fig. Photos of the lawn biofilm grown on 50% BSF/PBS with 1.5% agar after 48 hrs of static incubation (A).** Select areas of the plates without inoculated as a control (**B**) and with PAO1 (**C**) show the PAO1 biofilm has a brighter yellow/green color. The biomass generated is thinner than expected on LB agar so three plates were combined to compose one metabolomics sample.

**S3 Fig. Similar cell numbers as determined by CFUs were cultured from the biofilm and suspended culture samples for the untargeted metabolomics analysis.** Colony-forming units (CFUs)/mL for suspended cultures (blue) (**A**) (n=3) and CFUs/mL *×* cm^2^ for biofilm lawns (red) (**B**) (n=3) were measured by serial plate dilutions. When converted to CFU/culture (**C**) the cell count for suspended culture and biofilm was 2.5×10^11^ and 2.0×10^11^, respectively, which was not significantly different by unpaired, two-tailed *t*-test.

**S4 Fig. The two-dimensional score plot for PCA of the uninoculated BSF controls (green) (n=4), BSF suspended cultures (blue) (n=3), BSF lawn biofilms (red) (n=4), LB suspended cultures (teal) (n=9), and LB lawn biofilms (orange) (n=9).** PCA is based on the quantitation of metabolites peaks after normalization to total sum and pareto scaling showing clustering of sample cohorts with no overlap of the ellipses (ellipses represent 95% confidence intervals), displaying good separation between and repeatability within cohorts of samples. There is a clear separation between suspended culture and biofilm phenotypes regardless of media, but the separation is much greater between LB and BSF cultures.

**S1 Table. Metabolites identified and quantified from ^13^C-^1^H HSQC spectra of uninoculated BSF, PAO1 suspended in BSF, and lawn biofilm cultures grown in BSF.**

